# The *p*-coumaroyl arabinoxylan transferase *HvAT10* underlies natural variation in whole-grain cell wall phenolic acids in cultivated barley

**DOI:** 10.1101/2020.12.21.423816

**Authors:** Kelly Houston, Amy Learmonth, Ali Saleh Hassan, Jelle Lahnstein, Mark Looseley, Alan Little, Robbie Waugh, Rachel A Burton, Claire Halpin

## Abstract

Phenolic acids in cereal grains have important health-promoting properties and influence digestibility for industrial or agricultural uses. Here we identify alleles of a single BAHD *p-*coumaroyl arabinoxylan transferase gene, *HvAT10*, as responsible for the natural variation in cell wall-esterified *p*-coumaric and ferulic acid in whole grain of a collection of cultivated two-row spring barley genotypes. We show that *HvAT10* is rendered non-functional by a premature stop codon mutation in approximately half of the genotypes in our mapping panel. The causal mutation is virtually absent in wild and landrace germplasm suggesting an important function for grain arabinoxylan *p*-coumaroylation pre-domestication that is dispensable in modern agriculture. Intriguingly, we detected detrimental impacts of the mutated locus on barley grain quality traits. We propose that *HvAT10* could be a focus for future grain quality improvement or for manipulating phenolic acid content of wholegrain food products.

---

Phenolic acids in the cell walls of cereals limit digestibility^1^ when grain or biomass is used for animal feed or processed to biofuels and chemicals. They are also important dietary antioxidant, anti-inflammatory and anti-carcinogenic compounds and contribute to beer flavour and aroma^2,3^. The hydroxycinnamates, *p*-coumarate and ferulate (*p*CA and FA respectively), are the major phenolic acids in grasses. Both occur as decorations ester-linked to cell wall arabinoxylan. Lignin also has esterified *p*CA decorations but FA in lignin is incorporated directly into the growing polymer by ether linkages^4^. Besides its role as a lignin monomer, FA in the cell wall acts to cross-link arabinoxylans to each other and to lignin, and it is this cross-linking that may impede digestibility. The role of *p*CA in cell walls is less clear. In lignin, it may promote polymerisation of sinapyl alcohol monolignols^5^ and act as a termination unit^4^, but there are no clear theories about its role when attached to arabinoxylans. Given the importance of *p*CA and FA to plant health and the uses of cereal crops, there has been much recent interest in identifying genes that can be manipulated in transgenic plants to influence phenolic acid content^6-14^. Given current GM legislation in some countries it would be more appropriate for crop improvement to identify genes and alleles determining natural variation in *p*CA and FA that could be exploited immediately in contemporary plant breeding.

We quantified cell wall-esterified *p*CA and FA in the wholegrain of a replicated GWAS panel of 211 elite 2-row spring barley cultivars grown in a field polytunnel. We observed a 6-fold variation for esterified *p*CA (54 µg/g - 327 µg/g) and a greater than 2-fold variation in esterified FA (277 µg/g −748 µg/g) (Supplementary Fig. 1a,b, Supplementary Data 1, 2) with no correlation between FA and *p*CA levels (R^2^ = 0.04). A GWAS of this data using 43,834 SNP markers identified a single highly significant association for grain esterified *p*CA on chromosome 7H (-log10(p)=13.9; Fig. 1a, Supplementary Data and a co-locating peak for FA just below statistical significance (-log10(p)=3.9; Fig. 1b, Supplementary Data 3). Given the closeness of FA and *p*CA on the phenylpropanoid pathway we also conducted a GWAS using FA:*p*CA concentration ratios which provides internal data normalisation, reducing inherent variability in single compound measurements^15^. Mapping log[FA:*p*CA] values increased both the strength and significance of association with the locus (- log10(p)=19.4; Fig. 1c, Supplementary Fig. 2c, Supplementary Data 3), confirming a level of dependency between esterified FA and esterified *p*CA concentrations. GWAS on similar data from a semi-independent set of 128 greenhouse-grown barley genotypes identified the same associations (Supplementary Fig. 2a-c, Supplementary Data 3).

**Figure 1.**
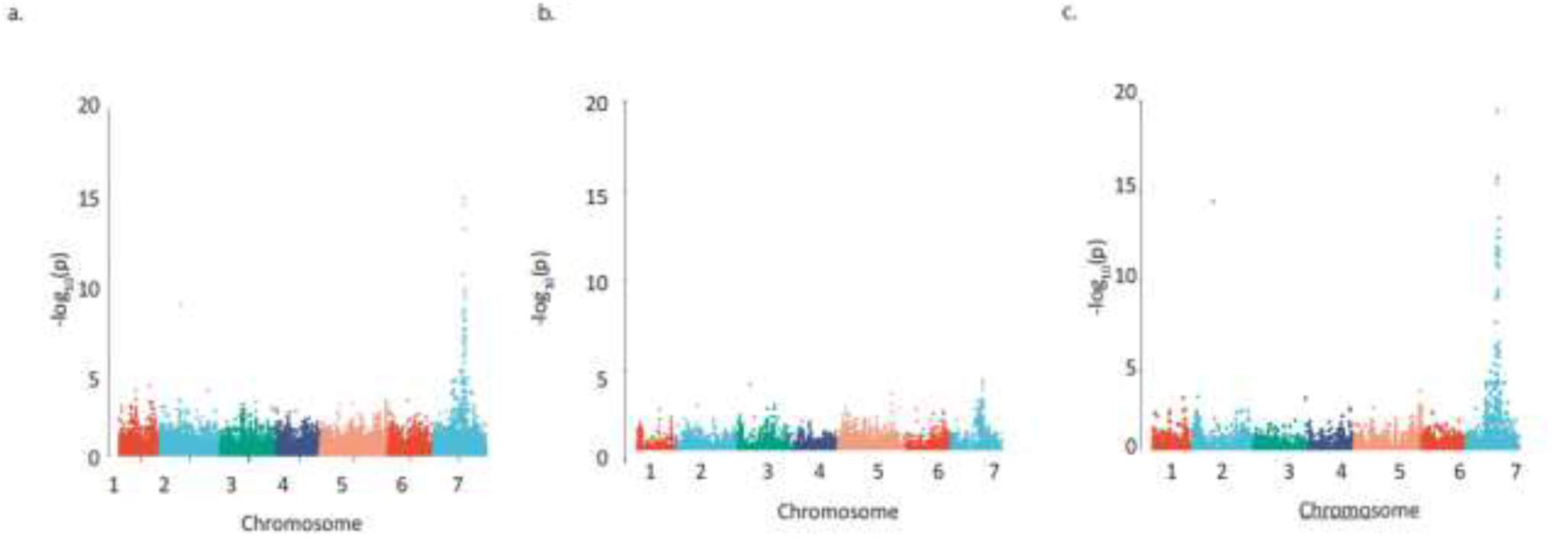
Detecting regions of the barley genome associated with grain phenolic acid content using a collection of 211 spring 2-row barleys. Manhattan plots of the GWAS of the phenolic acid content of wholegrain 2-row spring barley indicating regions of the genome associated with grain **a**. *p*- coumaric acid, **b**. ferulic acid content, **c**. using the a ratio of these two phenolic acids calculated by log[FA:p-Coumaric acid]. The –Log 10 (P-value) is shown on the Y axis, and the X axis shows the 7 barley chromosomes. The FDR threshold = −log 10(P)=6.02, plots use numerical order of markers on the physical map.

The entire region above the adjusted false discovery rate (FDR) threshold for the log[FA:*p*CA] values spanned a 65.7MB segment of chromosome 7H (459,131,547bp - 524,825,783bp) containing 347 high-confidence gene models. We surveyed this region for genes involved in phenolic acid or cell wall biosynthesis. This revealed several candidates including two cinnamyl alcohol dehydrogenases (*CAD*s), a caffeate-O-methyltransferase (*HvCOMT1*^16^) and three BAHD acyltransferases. Interrogation of an RNA-seq dataset for 16 barley tissues^17^ revealed that five of these six candidates exhibited moderate to low levels of expression across all surveyed tissues (Fig. 2a). However, the *BAHD* gene HORVU7Hr1G085100 stood out as being highly expressed in the hull lemma and palea where 80% of grain *p*CA is found^18^ (Fig. 2a, b). We then consulted a database of variant calls from a barley RNA-seq dataset that included 118 of our GWAS genotypes^19^. We observed no SNP variation in two of the candidate genes. Three had one SNP each; *COMT1* (HORVU7Hr1G082280) had a synonymous SNP, one *CAD* (HORVU7Hr1G079380) a SNP in the 3’ UTR and one *BAHD* (HORVU7Hr1G085390) a non-synonymous but rare SNP. None appeared likely to impair gene function. However, the *BAHD* HORVU7Hr1G085100 had 3 SNPs including one causing a premature stop codon leading to loss of a third of the protein sequence. BLASTp of the predicted full-length HORVU7Hr1G085100 protein sequence revealed it was 79% identical to rice *OsAT10* (LOC_Os06g39390.1), a gene functionally characterised as a *p*-coumaroyl CoA arabinoxylan transferase^7^. Critically, overexpression of *OsAT10* in rice dramatically increases cell wall-esterified *p*CA levels in leaves while concomitantly reducing the levels of esterified FA^7^. A maximum likelihood phylogenetic tree of *BAHD* gene sequences confirmed HORVU7Hr1G085100 as *HvAT10* (Fig. 2c) and another of our candidates, HORVU7Hr1G085390, as a possible *HvAT10* paralog with negligible expression in the tissues surveyed (Fig. 2a). The third *BAHD*, HORVU7Hr1G085060, is likely an AT8^7,13^.

**Figure 2.**
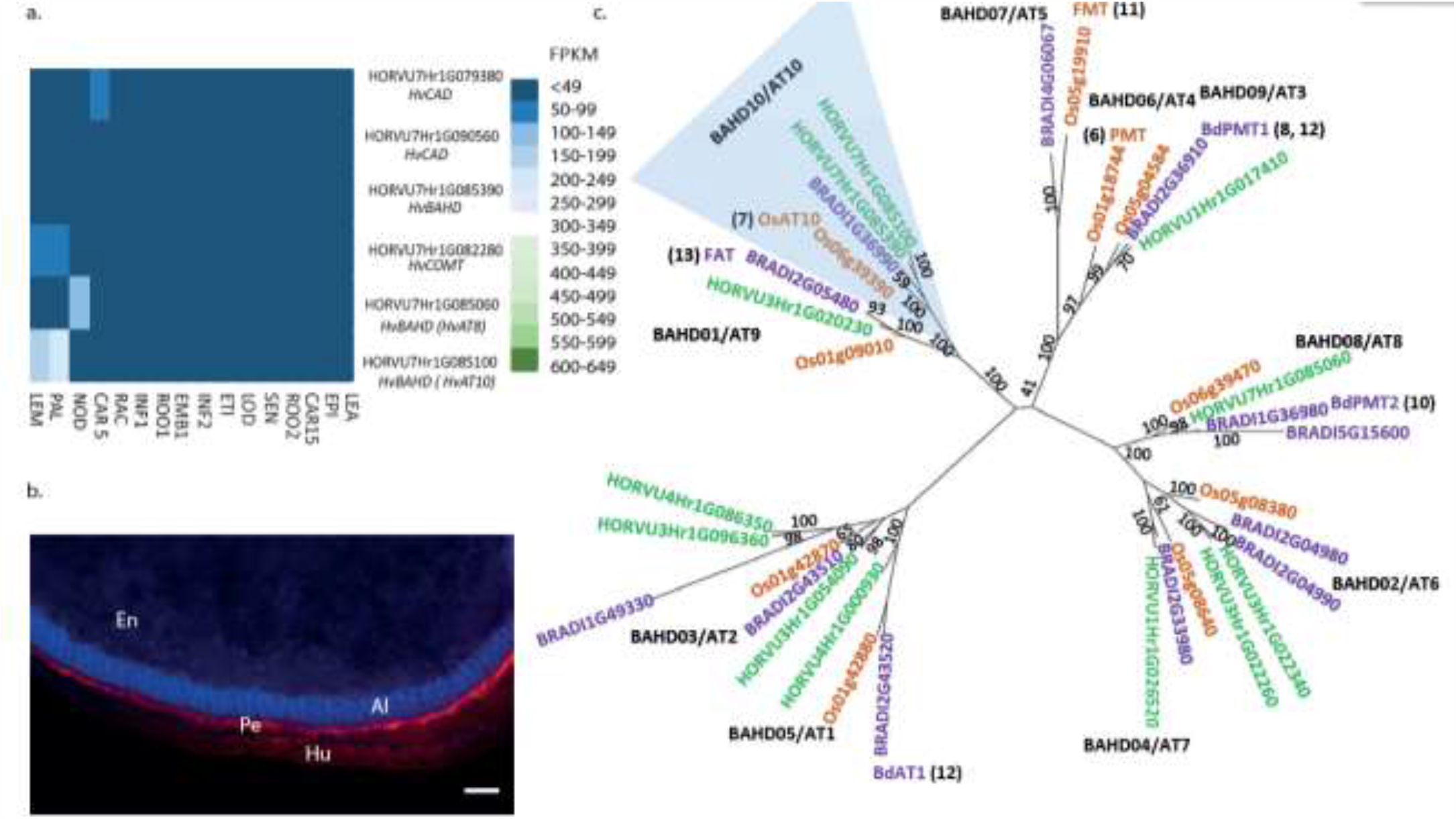
Putative candidates contributing to variation in grain *p*-coumaric (pCA) and Ferulic (FA) content from GWAS. **a**. Expression pattern for candidate genes under GWAS peak on 7H for grain *p*- coumaric and ferulic acid content in 16 different tissues/ developmental stages. Values are FPKM and a scale bar is provided. This expression data is derived from the publicly available RNAseq dataset BARLEX, https://apex.ipk-gatersleben.de/apex/f?p=284:39 **b**. Phenolic acid autofluorescence in whole grain sections. En: Endosperm, Al: Aleurone, Pe: Pericarp and Hu: Husk. Scale bar = 100μm **c**. Phylogenetic tree of the *BAHD* acyltransferases. A maximum-likelihood tree of the translation alignment of the coding sequences of group A and B *BAHD* genes from barley, rice and *Brachypodium*. Bootstrap support for branches is provided. Horvu numbers represent the barley gene models in green, BRADI represents *Brachypodium* in purple, and *Os* represents the rice genes in orange. The clade including LOC_OS06g39390 and HORVU7Hr1G085060 is highlighted in blue, *OsAT10* is indicated in green and the closest barley orthologue is marked in red. Where function of a gene model has been assigned the relevant reference is provided. Black text in bold indicates branch names, both BAHD and AT^13^.

To more accurately document polymorphisms in *HvAT10*, we PCR-sequenced the gene from 52 genotypes of the GWAS panel (Supplementary Data 1). Two nonsynonymous SNPs, one in each of *HvAT10*’s two exons (Fig. 3a), were in complete linkage disequilibrium across the 52 lines. A G/A SNP at 430bp translates to either a valine or isoleucine, substituting one non-polar, neutral amino acid for another, so unlikely to affect function. By contrast, a C/A SNP at 929bp produces either serine in the full length protein, or a premature stop codon that truncates the protein by 124 amino acids, removing the BAHD family conserved DFGWG motif (DVDYG in barley and other grasses) thought to be essential for catalysis^20-22^ (Fig. 3b). The *at10*^*STOP*^ mutation is therefore predicted to knock-out gene function. We designed a diagnostic Kompetitive Allele Specific PCR (KASP) assay to distinguish the two *HvAT10* alleles and genotyped all 212 cultivars in our GWAS population (Supplementary Table 1). Consistent with the hypothesis that *at10*^*STOP*^ is the causal variant underlying the log[FA:*p*CA] GWAS peak, no SNP scored higher than the KASP diagnostic when included in the GWAS although one, JHI-Hv50k-2016-488774, in complete LD scored equally highly. *HvAT10* had a minor allele frequency of 0.48 and appears to significantly influence levels of both *p*CA (*p*= 4.30e-19) and FA in grain (p=1.80e-11) with the median for *at10*^*STOP*^ genotypes being 28% lower for *p*CA (Fig. 3c) and 14% higher for FA (Fig. 3d) than those with the wildtype allele. Comparing the median log[FA:*p*CA] for *at10*^*STOP*^ cultivars (0.58) to the wildtype cultivar group (0.37) showed an even higher significant difference between the groups (*p*= 7.56e-50) (Fig. 3e, Supplementary Fig. 3).

**Figure 3.**
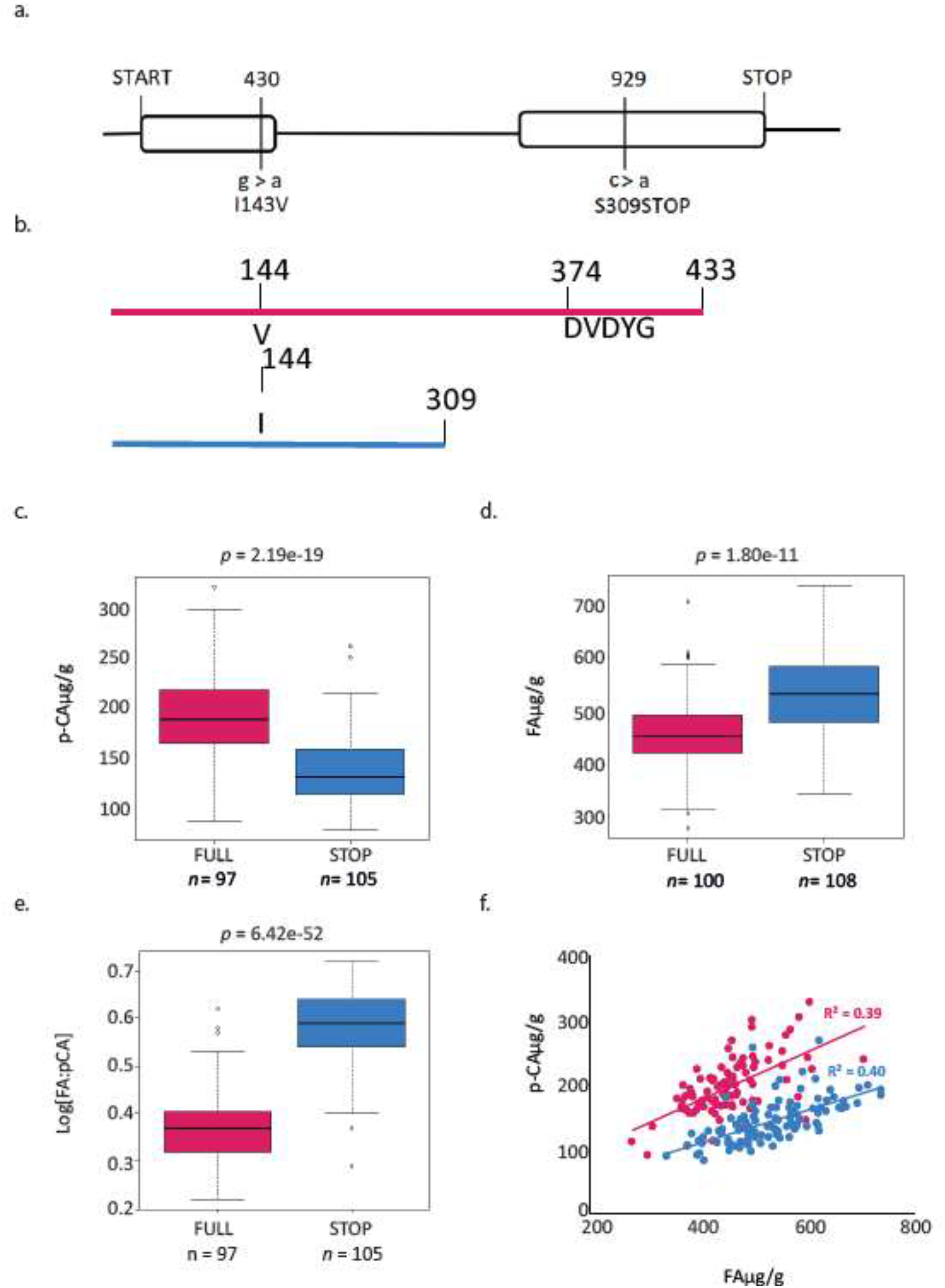
Gene and protein models for *HvAT10*. **a**. Gene model for *HvAT10* including location, and effect of SNPs detected from resequencing this gene in the 211 barley cultivars which have been assayed for *p*-coumaric and ferulic acid. The numbering above the gene model represent locations in the CDS which vary between these cultivars. The SNP, and the resulting change in the particular amino acid are indicated underneath the gene model. The full length of the gene is 2117bp (with a CDS of 1302bp) which translates to a protein of 435 amino acids as indicated. Protein model for translation of HvAT10. **b**. a Full length protein and **c**. when the premature stop codon is present this results in a truncated protein. Box plots demonstrate the effect of the SNP at 929bp within *HvAT10* where the grain of the 211 barley cultivars were quantified for **d**. *p*-coumaric acid levels and **e**. ferulic acid levels. **f**. Correlation between pCA and FA content based on *HvAT10* allele using 211 lines. The allele which results in full length version of HvAT10 are in pink, and the allele leading to a premature stop codon are coloured blue.

In contrast to our initial observation on the whole population, plotting grain esterified *p*CA against FA (Fig. 3f) within each allele group now reveals positive correlations, suggesting that although flux into phenolic acid biosynthesis may differ between cultivars, it co-ordinately affects both phenolic acids. The *at10*^*STOP*^ genotypes show approximately one-third less *p*CA than wildtype genotypes reflecting a deficiency of *p*CA on arabinoxylan in cultivars that lack a functional *p*-coumaroyl CoA arabinoxylan transferase. Nevertheless, two-thirds of cell wall esterified *p*CA remains since most *p*CA is associated with lignin^23,24^ through the action of other BAHD genes. The influence of *at10*^*STOP*^ on FA is evidenced by considering the 27 cultivars with grain esterified FA above 600 µg/g; 23 of these have the *at10*^*STOP*^ allele (Fig. 3f; Supplementary Data 1). This effect on FA might occur in several ways: *p*CA that cannot be esterified onto arabinoxylan could be methoxylated to produce FA thereby increasing FA pools for transfer onto arabinoxylan, or alternatively, *p*CA and FA may compete for transfer onto a shared acceptor (likely UDP-arabinose^12^) before incorporation into arabinoxylan such that loss of *p*CA transfer by *HvAT10* leaves more free acceptor for FA transfer. Either mechanism could explain how *at10*^*STOP*^ can indirectly increase grain cell wall esterified ferulate. An inverse interaction between levels of *p*CA and FA on arabinoxylan was also seen in transgenic rice^7^, switchgrass^26^, and *Setaria viridis*^13^ where BAHD expression was manipulated.

Intrigued by the prevalence of *at10*^*STOP*^ in 50% of our elite barley genepool we were curious about whether this had any ecological, evolutionary, or performance-related significance. To explore, we PCR-sequenced a collection of 114 georeferenced barley landraces and 76 wild barley (*Hordeum spontaneum*) genotypes^26^ across the *at10*^*STOP*^ polymorphism (Supplementary Data 1). We found *at10*^*STOP*^ to be extremely rare, present in three of 114 landraces and absent in all 76 wild genotypes (Supplementary Fig. 4a, Supplementary Data 1). The three landraces show a clear pattern of identity by descent, clustering in the same clade of the dendrogram (Supplementary Fig. 4a). We interpret these data as suggesting strong selection against the premature stop codon in wild germplasm and that *at10*^*STOP*^ was a post-domestication mutation that under cultivation has no pronounced negative effects on fitness.

Several possibilities could explain enrichment of *at10*^*STOP*^ in the cultivated genepool. To explore, we first calculated genome wide F_ST_ by locus using two groups based on the *HvAT10* allele. HORVU7Hr1G084140 (a Serine/threonine-protein kinase not expressed in the lemma or palea) also had an F_ST_ of 1.0, and three other genes an F_ST_ above 0.875 (Supplementary Fig. 5a,b Supplementary Data 4). Based on their functional annotations and gene expression patterns (Supplementary Data 4, Supplementary Fig. 5c) we observed no obvious reason for these to be under strong selection and responsible for enhancing the frequency of *at10*^*STOP*^ via extended LD.

Next, due to the exclusive expression of *HvAT10* in the lemma and palea, we measured a series of grain morphometric traits across our panel. We found that, on average, grain from the *at10*^*STOP*^ genotypes had significantly reduced grain width compared to cultivars with the wildtype allele (Table 1) suggesting a potential role for arabinoxylan-esterified phenolic acids in modifying grain shape. Xu *et al*^27^ previously identified a QTL hotspot on chromosome 7H for traits including grain area, and grain width. The eight 9K iSelect markers defining this QTL can be positioned on the current physical map at 482-500MB on 7H, corresponding to the location of *HvAT10*. Wang *et al* ^28^ also identified a QTL for grain length:width, grain perimeter, and grain roundness at the same location.

**Table 1.** T-test results for comparisons between *HvAT10* alleles. For grain area, length and width data are available in Supplementary Data 1 and analysis was carried out using BLUPS derived from 2 – years’ worth of samples. Data used for comparison of hot water extract, germinative energy, fermentable extract, diastatic power, wort viscosity and friability between *HvAT10* alleles are published ^29^.

|  | Grain Area | Grain Length (mm) | Grain Width (mm) | Hot water extract (g/Kg) | Germinative energy 4 ml (%) | Fermentable extract | Diastatic power loB | Wort viscosity (mPa/s) | Friability (%) |
| --- | --- | --- | --- | --- | --- | --- | --- | --- | --- |
| *at10 STOP* | 27.67 | 9.02 | 3.92 | 309.44 | 97.34 | 70.72 | 100.77 | 1.5 | 87.97 |
| WT | 28.16 | 9.09 | 3.97 | 311.23 | 97.4 | 70.88 | 106.07 | 1.48 | 89.95 |
| *p* value | 0.0214* | 0.152 | 0.0009*** | 0.0005*** | 0.0206* | 0.0203* | 0.0282* | 0.0009*** | 0.0123* |

Prompted by these observations and the prevalence of registered UK barley varieties in our panel, we then explored grain parameters recorded in an extensive historical dataset from the UK’s National and Recommended Lists trials 1988-2016^29^. Different grain quality phenotypes were available for up to 106 of our cultivars. Group comparisons of WT and *at10*^*STOP*^genotypes revealed surprising differences for hot water extract, diastatic power, germinative energy in 4ml, and wort viscosity (Table 1). In all cases, the group of *at10*^*STOP*^ cultivars had poorer quality, offering no evidence of positive selection during breeding. The variation associated with the *HvAT10* locus is however highly significant and of potential interest for optimising grain quality traits (Table 1). Finally, to understand more about the origin of the *at10*^*STOP*^ in elite germplasm, we investigated its occurrence in the pedigree of our GWAS population. The earliest cultivar with *at10*^*STOP*^ is *cv*. Kenia (cross between the Swedish landrace Gull and Danish landrace Binder) released in 1931 and subsequently introduced into NW European breeding programmes. Despite smaller grain and slightly poorer malting properties compared to its contemporary UK varieties, it established a long-standing position as a parent for further crop improvement due to its short stiff straw, earliness and high yield^30^. Several decades later, *at10*^*STOP*^-containing derivatives of Kenia, such as *cv*. Delta (National list 1959), were still being used as parents in our pedigree chart.

Taken together, we conclude that the continued prevalence of Kenia-derived germplasm may go some way to explaining the frequency of the *at10*^*STOP*^ allele in our population. While this may simply be a straightforward genetic legacy of historical barley breeding, our data suggests that purging this mutation could assist the development of superior quality barley varieties. Conversely, much research has focussed on the beneficial bioactivity of ferulate in the diet and the *at10*^*STOP*^ allele could enable breeding for increased ferulate in wholegrain products.

## Methods

### Plant material and growth conditions

Two populations of 2-row spring type barley were used to carry out the GWAS^31^. The first population includes 211 elite lines grown in a polytunnel under field conditions in Dundee, Scotland. For each line, 5 whole grains were ground to a fine powder using a ball mill (Mixer Mill MM400; Retsch Haan Germany) and stored in dry conditions until the HPLC analysis. The second population which was used for verification of the results of the analysis of the first subpopulation includes 128 elite lines grown in a glasshouse compartment in a mix of clay-loam and cocopeat (50:50 v/v) at daytime and night time temperatures of 22°C and 15°C respectively in The Plant Accelerator, Adelaide, Australia. As described previously, the collection of germplasm these populations are sampled from has minimum population structure while maintaining as much genetic diversity as possible^32^. Mature grains were stored until phenolic acid content analysis.

### Genotyping of SNP markers

All lines were genotyped using the 50K iSelect SNP genotyping platform described previously^33^. Prior to marker-trait association analysis, all markers with a minimum allele frequency of <5% and markers with missing data >5% were excluded from the analysis.

### Phenotyping for cell wall-bound phenolic acids

A ∼ 20 mg amount of wholegrain barley was used per sample. *Trans*-ferulic and *trans*-*p*-coumaric acid standards were purchased from SIGMA Aldrich (Castle Hill NSW, Australia). Standards were prepared at 62.5 µm, 250 µm and 1000 µm by dissolving the appropriate amount of powder in 50% methanol. Extraction of cell wall esterified phenolic acids was carried out following the methods described by ^34,35^ with the following modifications. Samples were washed twice with 500 µl 80% ethyl alcohol, with shaking for 10 minutes at room temperature to remove free phenolic acids. To release total cell wall esterified phenolic acids, alkaline treatment was carried out by adding 600 µl 2M NaOH to the pellet. Samples were incubated on a rotary rack under nitrogen for 20 h in the dark at room temperature. Samples were centrifuged at 15000 × g for 15 minutes at room temperature, after which the supernatant was collected, acidified by adding 110 µl concentrated HCL and extracted three times with 1 mL ethyl acetate. Following each extraction, samples were centrifuged at 5000 × g for 7 minutes and the organic solution was collected. Extracts were combined, evaporated to dryness in a rotary evaporator and dissolved in 100 µl of 50 % methanol prior to injecting 40 µl into the HPLC column. For each sample two technical replicates were applied.

### HPLC conditions

An Agilent Technologies 1260 Infinity HPLC equipped with a Diode Array detector was used. Samples were analysed on an Agilent Poroshell 120 SB-C18 3.0×100mm 2.7- micron column kept at 30 C°. Eluents were A (0.5mM trifluoroacetic acid) and B (0.5mM trifluoroacetic acid, 40% methanol, 40% acetonitrile, 10% water). Starting conditions were 85% A and 15% B. Flow rate was 0. 7 mL/min. Eluting gradients were as follow; min 0-10: 15% to 55% B, min 11-12: column washed with 100% B, min 13 back to the starting condition (85% A and 15% B). Detection was carried out at 280 nm and spectral data was collected from 200 to 400 nm when required. Ferulic and *p*-coumaric acid peaks were identified by comparing retention times and spectra to their corresponding standards. The area under the peaks was quantified at 280 nm for *trans* forms.

### GWAS analysis of grain alkaline extractable pCA and FA and FA:pCA ratio

Marker- trait association analysis was carried out using R 2.15.3 (www.R-project.org) and performed with a compressed mixed linear model^36^ implemented in the GAPIT R package^37^. For phenotype values, the mean values of the barley wholegrain total alkaline extractable *trans*-ferulic and *trans*- *p*- coumaric acid (w/w) were used. To identify genes within intervals associated with our trait we used ^17^. We also used the ratio of FA:pCA as a trait in our GWAS analysis. The ratio between the two compounds was log transformed i.e. log(FA:pCA) to provide a more normally distributed dataset. When using ratios in GWAS, a significant increase in the *p*-gain statistic^15^ (a comparison between the lowest -log10(p) values of the individual compounds and the -log10(p) value of the ratio) indicates that ratios carry more information than the corresponding metabolite concentrations alone. A significant p-gain identifies a biologically meaningful association between the individual compounds. We used B/(2*α) to derive a critical value of 3.42×10^5^ for the FDR-adjusted p-gain, where α is the level of significance (0.05) and B the number of tested metabolite pairs^15^. Therefore, as we tested two traits our threshold was 2 ൿ 10^1^ and our p-gain was above this threshold.

To identify local blocks of LD, facilitating a more precise delimitation of QTL regions Linkage disequilibrium (LD) was calculated across the genome between pairs of markers using a sliding window of 500 markers and a threshold of R^2^<0.2 using Tassel v 5 ^38^. We anchored markers that passed FDR and represented initial borders of the QTL on 7H to the physical map and then expanded this region using local LD derived from genome wide LD analysis as described above. When the GWAS had not resulted in an association that passed the FDR we used the arbitrary threshold of - LOG10(P) to define the initial border. The SNP with the highest LOD score was used to represent the QTL. After identification and Sanger sequencing of the candidate gene *HvAT10* the GWAS was repeated including the allele present at the S309Stop as an additional marker.

### Bioinformatics and gene identification

We used BARLEX^17^ to identify gene models present with the QTL defined by our analysis and their expression profile based on RNAseq data in 16 different tissues/ developmental stages

### Phylogenetic analysis of barley BAHD acyltransferases

Coding sequences of all BAHD acyltransferases with the PFAM domain PF02458 from rice, barley and *Brachypodium* were downloaded from the Ensembl Plants database (http://plants.ensembl.org/). Sequences were aligned using the MUSCLE alignment function^39^available in the Geneious 9.1.4 (https://www.geneious.com). The translation alignment option was used. A neighbour-joining tree was produced from the alignment. Barley genes within group A and B clades were identified, realigned with their rice and *Brachypodium* orthologs and a maximum likelihood tree was produced from the translation alignment of the sequences. The following settings were applied: substitution model: General-Time-Reversible (GTR), branch support: bootstrap, number of bootstrap: 1000.

### Resequencing and genotyping of *HvAT10* in the main and supplemental set

Aligning the translation of AK376450 to Os06g39390 allowed the identification of the putative genomic sequence of *HvAT10*. We designed four pairs of primers, details of sequences and reaction conditions are in Supplementary Data 5, to amplify the full length CDS using reaction volumes, reagents, and conditions as described in^40^. To facilitate quick and efficient genotyping of large numbers of cultivars we subsequently designed a KASP genotyping assay to a SNP at 430bp in *HvAT10* (Supplementary Data 5). Reactions were performed in an 8.1 µL reaction volume, with 3 µL H2O, 1 µL DNA (20ng/µl), 4 µL KASP genotyping master mix, and 0.11 µL of the KASP assay.

Box plots to demonstrate the contribution of the SNP at 436bp in *HvAT10* to variation in grain pCA and FA content were produced using R 2.15.3 (www.R-project.org). To test for identity by descent of the *HvAT10* allele within the set of accessions using for the GWAS a dendrogram was constructed using maximum likelihood using the genotypic data from the 9k-select array^32^ in MEGA7^41^ with default settings except for including bootstrapping and visualised in FigTree (v.1.4.4) http://tree.bio.ed.ac.uk/software/figtree/.

### Characterisation of diversity of *HvAT10* in *H. spontaneum* from the fertile crescent and barley landraces

DNA was extracted as described above from 76 *H. spontaneum* and 114 barley landraces from^26^. The S309Stop SNP was PCR amplified and Sanger sequenced with primer pair 5 using conditions described above. A dendrogram was constructed using maximum likelihood using 4000 exome capture derived SNPs from^26^ in MEGA7^41^ with default settings except for including bootstrapping and visualised in FigTree (v.1.4.4) http://tree.bio.ed.ac.uk/software/figtree/.

### Genome wide F_ST_ analysis

The fixation index (F_ST_) is a measure of genetic differentiation between groups of individuals. Genome wide F_ST_ was calculated by locus using GenAlEx 6.502^42,43^ after dividing the accessions into two populations based on their HvAT10 allele using all informative 50K iSelect markers.

### Phenotypic analysis of cultivars with wildtype vs *at10*^*STOP*^ allele

We characterised mature grain morphology using from plants grown in a polytunnel under field conditions in Dundee, Scotland as described above, over two years (2010 and 2011). Grain area, width and length were quantified using the MARVIN Seed Analyzer (GTA Sensorik GmbH, 2013). BLUPs calculated from this data using R 2.15.3 (www.R-project.org) were used in subsequent comparisons between allelic groups.

## Data availability

All sequences of *HvAT10* generated in this study are available from NCBI, accession numbers are provided in **Supplementary Table 1**.

## Acknowledgements

KH and RW acknowledge support from the Rural & Environment Science & Analytical Services Division of the Scottish Government and BBSRC (BB/J014869/1, BB/L026317/1). CH and RW are grateful for funding from BBSRC (BB/N023455/1 and BB/G016232/1). Amy Learmonth was supported by BBSRC grant BB/J01446X/1. ML and KH acknowledge funding from BB/K008188/1. We would also like to acknowledge technical support from Malcolm Macaulay and advice on analysis from Helen Oakey. RB, ASH, and Alan Little acknowledge funding provided by the Australian Research Council Centre of Excellence in Plant Cell Walls CE110001007.

## Author contributions

RW, KH, RB, CH, Alan Little, designed experiments. KH, ASH, JL, Amy Learmonth, carried out experiments. KH, ASH, Amy Learmonth, ML, Alan Little, JL analysed data. The manuscript was written by CH, KH, RW, RB, Amy Learmonth, ASH with contributions from all other authors.

## Ethics declarations

The authors declare no competing interests.

## Supplementary Information

**Supplementary Data 1. Phenolic acid and genetic data for all cultivars included in this study**. *p*- coumaric and ferulic acid content, KASP data and NCBI number for those lines that where sequenced for *HvAT10* is included.

**Supplementary Data 2. Summary of number of accessions used for each GWAS**. A total of 211 accessions were included in the main dataset but data for both phenolic acids is not available for all lines, therefore the number of individuals included in different analysis varies. Includes number of accessions for the GWAS presented in the main and supplementary analysis for both individual trait and the ratio analysis. Number of individuals with each allele of *HvAT10* based on genotyping of A430G is also included.

**Supplementary Data 3. Details of QTL identified on 7H for all analysis carried out**. Physical location, LOD score, and 50k iSelect marker with the highest LOD score are provided. * indicates that analysis passed the FDR threshold of -log10(p)=6.1.

**Supplementary Data 4. Gene models containing SNPs that have an F**_**ST**_**>0.875 when F**_**ST**_ **analysis carried out based on *HvAT10* allele**. This table includes 50k iSelect marker name, the chromosome the marker is located on, gene model and annotation based on Morex v1 Gene Models (2016).

**Supplementary Data 5. Details of primers and genotyping assays used in this study**. This includes details of primers for Sanger sequencing and KASP genotyping assay sequence for *HvAT10*.

**Supplementary Figure 1. Phenolic acid content of wholegrain flour from 211 2-row spring barleys linea. a**. *p*CA and **b**. ferulic acid content. Values represent the mean for FA and pCA expressed as w/w. Error bars represent standard deviation of the replicates.

**Supplementary Figure 2. Manhattan plots of the GWAS of the phenolic acid content of wholegrain flour from 128 2-row spring barley lines indicating regions of the genome associated with grain phenolic acid content**. Manhattan plots of the GWAS of the phenolic acid content of wholegrain 2- row spring barley indicating regions of the genome associated with grain **a**. *p*-coumaric acid, **b**. ferulic acid content and **c**. log[FA:p-Coumaric acid]. The –Log 10 (P-value) is shown on the Y axis, and the X axis shows the 7 barley chromosomes. FDR threshold?=?−log 10(P)=6.02, plots use numerical order of markers on the physical map.

**Supplementary Figure 3. Distribution of ratio between two phenolic acids quantified in the grain of 211 spring 2 row barleys lines and used to carry out GWAS**. The ratio was calculated as log[FA:p-Coumaric acid]. Accessions containing the allele which results in a full length version of *HvAT10* are in pink, and accessions containing the allele leading to a premature stop codon are coloured blue.

**Supplementary Figure 4. Distribution of the *HvAT10* premature stop codon in *H. vulgare* landraces and cultivated barley lines. a**. A dendrogram of 114 *H. vulgare* landraces constructed using a selection of SNPs with a genome-wide distribution with maximum likelihood methods. **b**. A dendrogram of cultivated barley germplasm using a selection of SNPs with a genome-wide distribution using maximum likelihood methods. Accessions containing the allele which results in full length version of HvAT10 are in pink, and accessions containing the allele leading to a premature stop codon are coloured blue.

**Supplementary Figure 4. F**_**ST**_ **analysis based on *HvAT10*. a**. Plot displaying genome wide F_ST_ with F_ST_ index provided on the Y axis, an F_ST_ of 1 indicating a complete fixation of each allele within the two subpopulations determined by their allele of *HvAT10*. **b**. Just F_ST_ of markers at 7H. Red box indicates location of the centromere. Two SNPs whose location overlap on this plot, including one in *HvAT10*, have an F_ST_ of 1.0. Note shape of peak appears different in **a**. and **b**. due to the difference in scale of the plots. **c**. RNAseq data for genes with F_ST_>0.875 from 16 different tissues/ developmental stages. Values are FPKM and a scale bar is provided. This expression data is derived from the publicly available RNAseq dataset BARLEX, https://apex.ipk-gatersleben.de/apex/f?p=284:39

