## Supplementary figures and images for "The *p*-coumaroyl arabinoxylan transferase *HvAT10* underlies natural variation in whole-grain cell wall phenolic acids in cultivated barley"

### Supplemental Figure 1

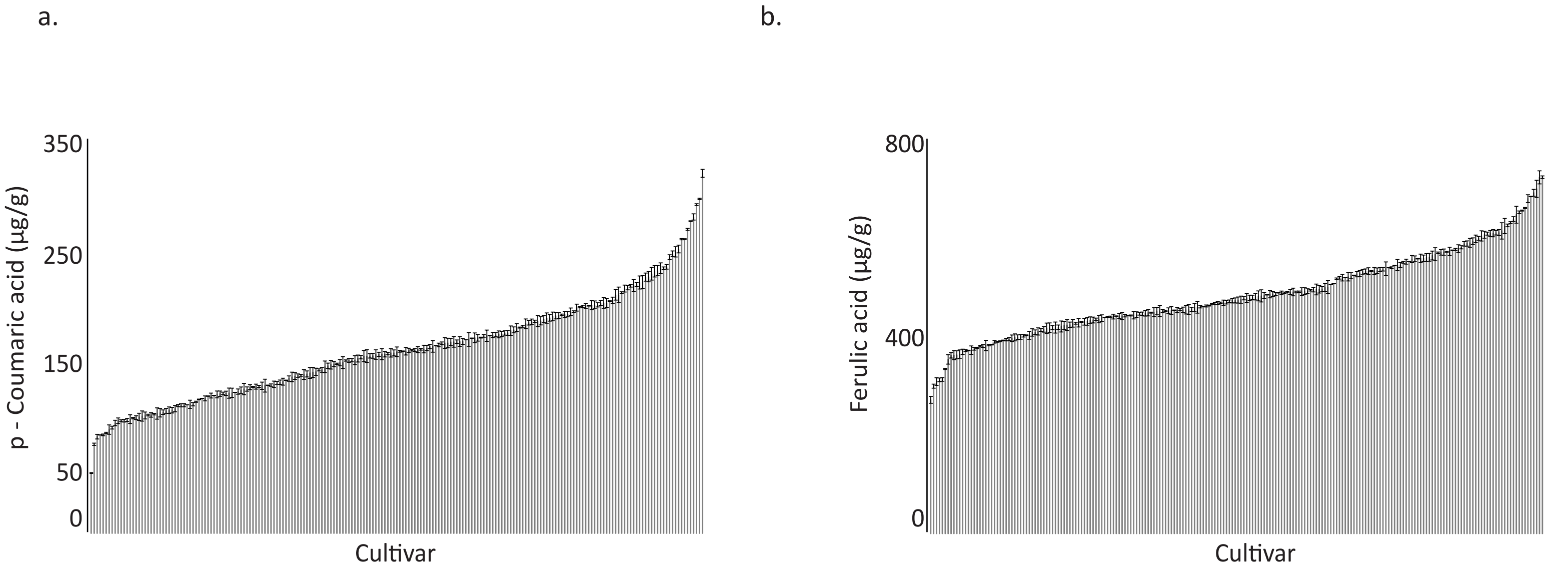

### Supplemental Figure 2

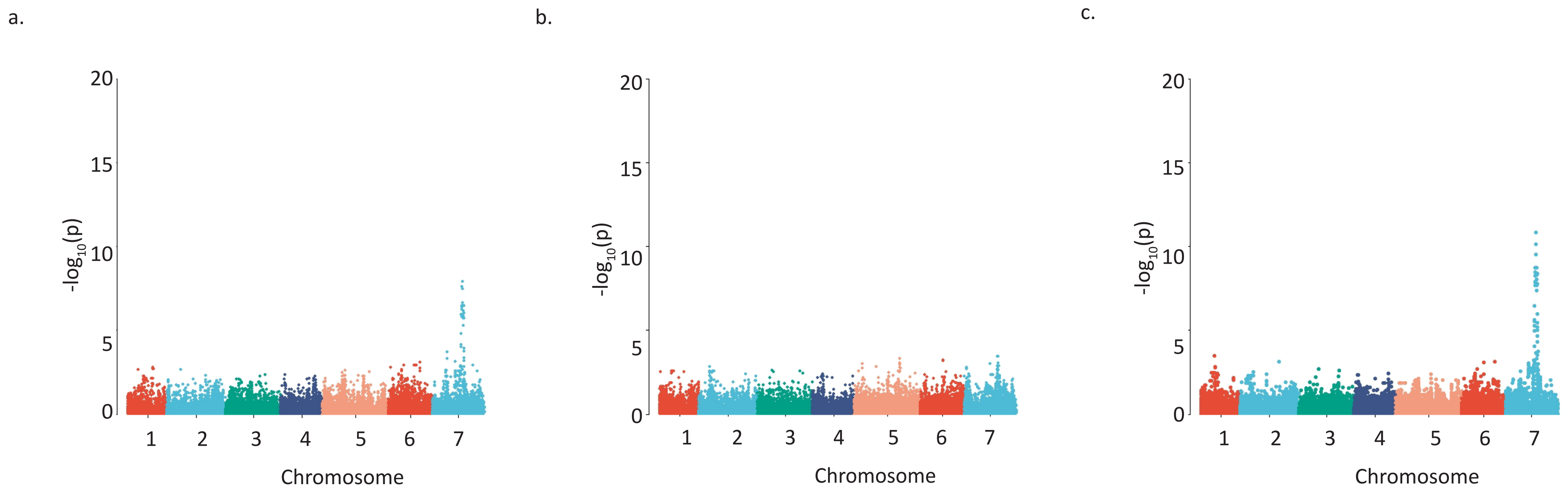

### Supplemental Figure 3

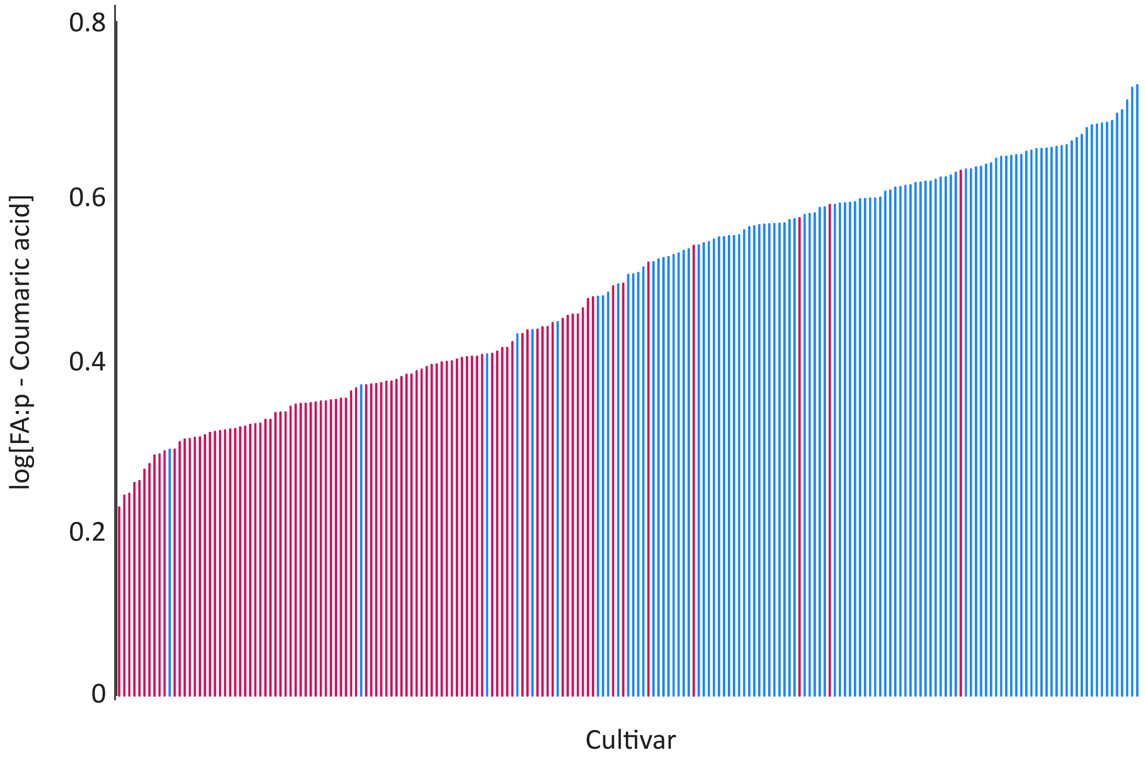

### Supplemental Figure 4

a.

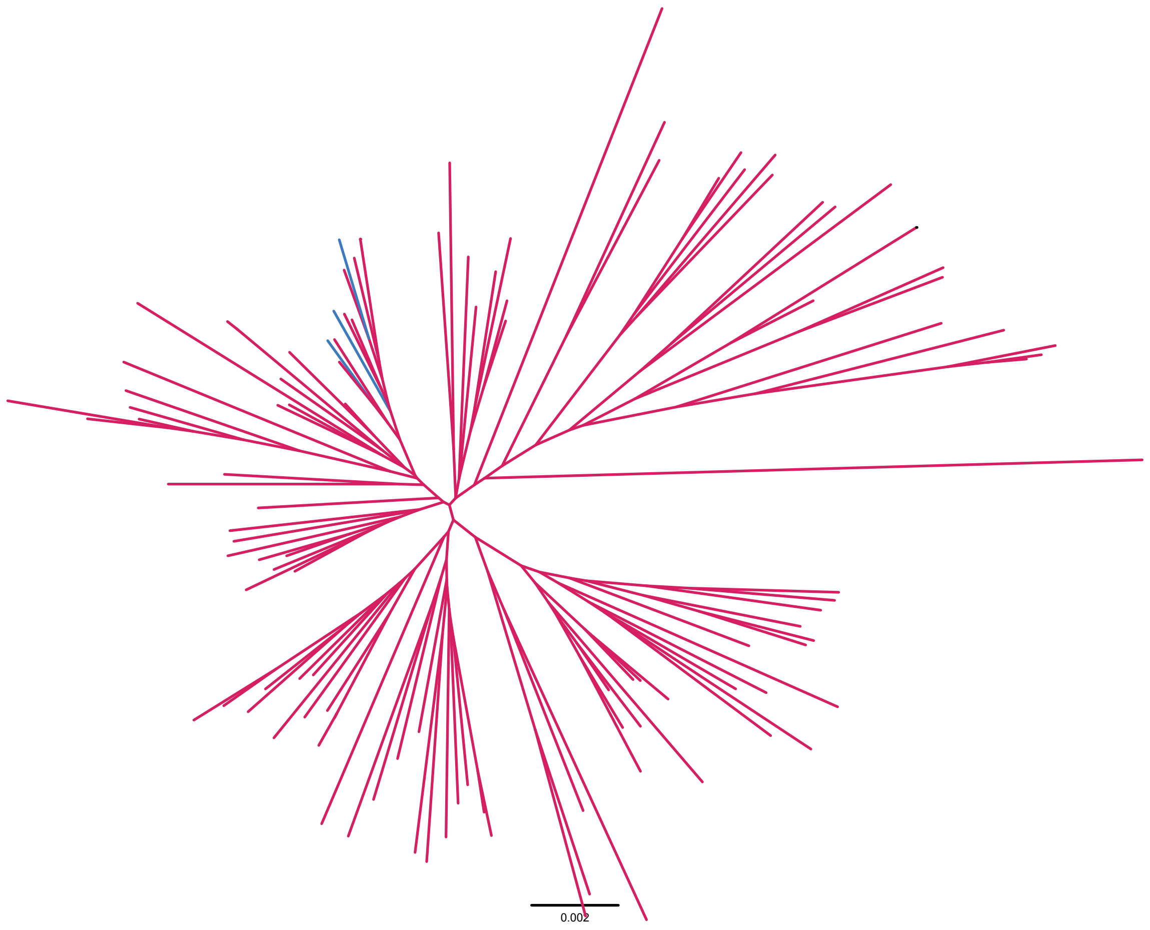


b.

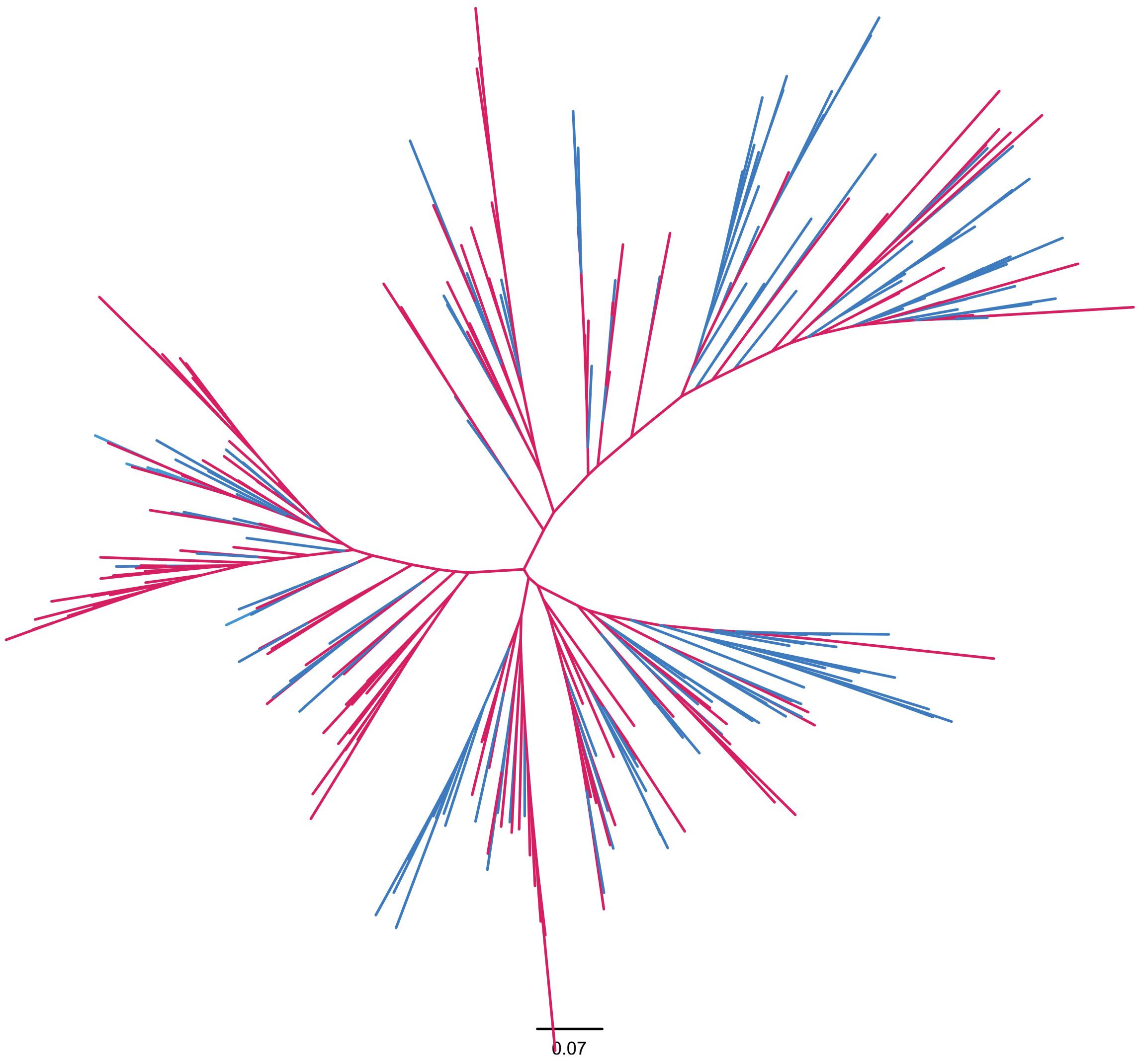

### Supplemental Figure 5

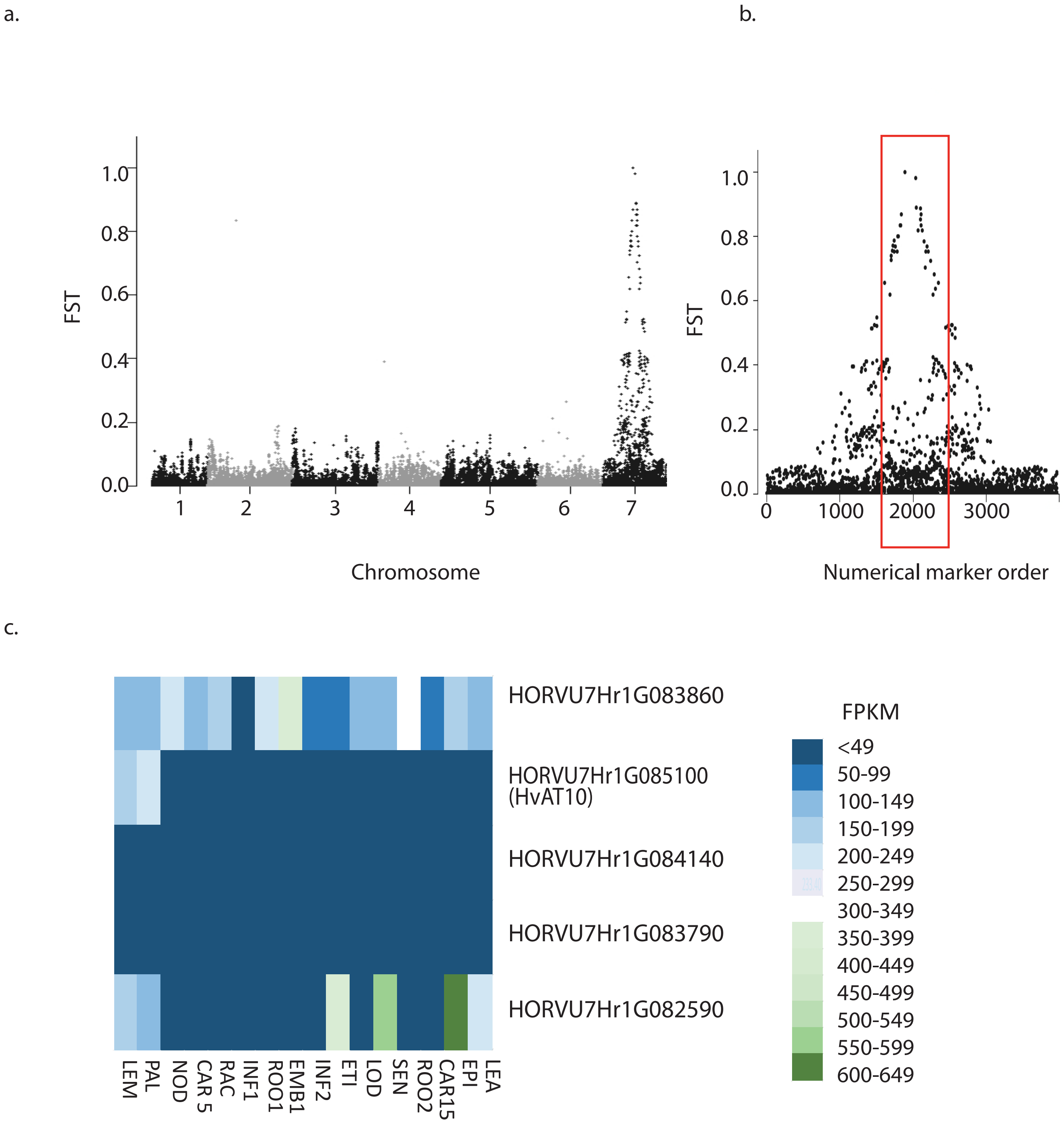
